# Global variation in the SARS-CoV-2 proteome reveals the mutational hotspots in the drug and vaccine candidates

**DOI:** 10.1101/2020.07.31.230987

**Authors:** L Ponoop Prasad Patro, Chakkarai Sathyaseelan, Patil Pranita Uttamrao, Thenmalarchelvi Rathinavelan

**Affiliations:** Department of Biotechnology, Indian Institute of Technology Hyderabad Kandi, Telangana-502285

**Keywords:** SARS-CoV-2 viromics, proteome analysis, mutational hotspot, mutational susceptibility, phyloproteomics, drug target, vaccine, diagnostics

## Abstract

To accelerate the drug and vaccine development against the severe acute respiratory syndrome virus 2 (SARS-CoV-2), a comparative analysis of SARS-CoV-2 proteome has been performed in two phases by considering manually curated 31389 whole genome sequences from 84 countries. Among the 9 mutations that occur at a high significance (T85I-NPS2, L37F-NSP6, P323L-NSP12, D614G-spike, Q57H-ORF3a, G251V-ORF3a, L84S-ORF8, R203K-nucleocapsid and G204R-nucleocapsid), R203K-nucleocapsid and G204R-nucleocapsid are co-occurring (dependent) mutations and P323L-NSP12 and D614G-spike often appear simultaneously. Other notable variations that appear with a moderate to low significance are, M85-NSP1 deletion, D268-NSP2 deletion, 112 amino acids deletion in ORF8, a phenylalanine insertion amidst F34-F36 (NSP6) and several co-existing (dependent) substitution/deletion (I559V & P585S in NSP2, P504L & Y541C in NSP13, G82 & H83 deletions in NSP1 and K141, S142 & F143 deletions in NSP2) mutations. P323L-NSP12, D614G-spike, L37F-NSP6, L84S-ORF8 and the sequences deficient of the high significant mutations have led to 4 major SARS-CoV-2 clades. The top 5 countries bearing all the high significant and majority of the moderate significant mutations are: USA, England, Wales, Australia and Scotland. Further, the majority of the significant mutations have evolved in the first phase and have already transmitted around the globe indicating the positive selection pressure. Among the 26 SARS-CoV-2 proteins, nucleocapsid protein, ORF3a, ORF8, RNA dependent RNA polymerase and spike exhibit a higher heterogeneity compared with the rest of the proteins. However, NSP9, NSP10, NSP8, the envelope protein and NSP4 are highly resistant to mutations and can be exploited for drug/vaccine development.

## Introduction

The severe acute respiratory syndrome virus 2 (SARS-CoV-2) pandemic outbreak has led to 1,36,99,958 and 5,91,350 global infected cases and deaths respectively as of July 15, 2020 due to the lack of proper diagnostics and therapeutics. Treating the SARS-CoV-2 pandemic is the biggest challenge the world currently faces and finding an effective vaccine or drug molecule is of paramount importance now. Several attempts are being made in drug repurposing against the viral target to treat SARS-CoV-2 infection^1–4^. Although several vaccines are under pipeline, their effectiveness is still unpredictable^3,5,6^. A comparative analysis of SARS-CoV-2 clinical genome/proteome sequences from different geographical location would provide information on the mutational rate and would facilitate the understanding of the viral evolution, its implication in the molecular mechanisms and pathology of the disease in different populations.

Although SARS-CoV-2 is expected to have slow mutation rate due to its sophisticated proof reading mechanism^7^, it can incorporate changes in the genome ^8–10^ to escape from the host immune defenses. A comparative analysis of 31389 SARS-CoV-2 whole genome sequences obtained from the clinical samples (deposited in NCBI and GISAID^11,12^ till May 17, 2020) from 84 countries are exploited here to accelerate the identification of universal vaccine candidates and drug targets. A country wise local proteome repository has been created and subjected to multiple sequence alignment to analyze the rate of mutation in two different phases (till April 17, 2020 and during April 18-May17, 2020).

The SARS-CoV-2 genome has 10 open reading frames (ORFs) flanked by 5’UTR and 3’UTR ^13,14^. Upon entering into the host either by direct membrane-envelope fusion or receptor-mediated endocytosis, the ORF1 gets translated into two large polyproteins (PP1a and PP1ab) with the utilization of the host translational machinery. PP1a consists of non-structural proteins (NSPs) 1 to 11, whereas, the PP1ab comprises of NSPs 1 to 16 (except NSP11) from the genomic RNA. The matured NSP1 to NSP16 are produced with the help of two viral encoded cysteine proteases, namely, a papain-like protease (PL^pro^) (proteolysis NSP1-3) and a main protease (M^pro^ or 3CL^pro^) (proteolysis NSPs 4 to 16). The NSPs7-16 act as the replication machinery that enables the synthesis of the viral genome. The subgenomic RNAs from S,ORF3a,E,M,ORF6,ORF7a,ORF7b,ORF8,N and ORF10 are involved in the translation of virion structural and accessory proteins. While the proteins synthesized by ORF3a,ORF6,ORF7a,ORF7b,ORF8 and ORF10 are designated as accessory proteins, the proteins from S,E,M and N are designated as structural proteins ^13^. ORF10 is utilized in the diagnostics to differentiate SARS-CoV from SARS-CoV-2 as it is unique in the latter^15,16^.

SARS-CoV-2 proteome analysis indicate that there are 778, 32 and 9 low (LS), moderate (MS) and high (HS) significant mutations respectively common to both the phases which led to 4 major clades. Among which, D614G-spike and P323L-NSP12 are found in ~75% of the total number of sequences in phase 2. Interestingly, R203K and G204R present in nucleoprotein are found to be co-occurring (dependent) mutations. The study also reports the presence of several deletions and co-occurring mutations with a moderate significance. As envelope protein, NSP4, NSP9 and NSP10 are less heterogenous compared with the rest of the proteins, they can be good vaccine/drug targets. Thus, the results presented here would help in identifying the potential therapeutic targets. It may further have an impact on correlating the mutations with the viral evolution, virulence and fatality.

## Results

31389 sequences have been considered to analyze SARS-CoV-2 proteome diversity across 84 countries and 46 countries have data in both the phases. While, 22 countries have data only in the first phase, 16 countries have data only in the second phase.

### Mutations in the proteins encoded by ORF1ab

#### Evolution of new lineages with K141-S142-F143 and G82-H83 co-deletions in NSP1

NSP1 is a 180 amino acid length protein which participates in host mRNA degradation and immune inactivation^17^. Out of the 84 countries, 29 countries (England, Wales, Scotland, France, Luxembourg, Iceland, Netherlands, Portugal, Spain, Belgium, Norway, Austria, Ireland, Denmark, Greece, Sweden, Switzerland, Japan, Russia, Saudi Arabia, Singapore, South Africa, India, Israel, UAE, USA, Canada, Australia and New Zealand) have 27 unique and significant mutations (in 23 positions) at least in one of the phases **(Figure 1A, B)**. There are 18 mutations common to both the phases, for which, the most of the mutations exhibit an increase in the mutation rate in the second phase compared with the first phase (**Figure 1**) (see Methods). Notably, R24C, G82E, A117T and G150C that are significant (mutation rate of about 0.04%) in the first phase has become insignificant in the second phase (**Figure 1)**. In contrast, H45Y, A76T, V84H, M85V, A90V and D139Y that are either absent or insignificant in the first phase have become significant (mutation rate ranges between 0.04% and 0.12%) in the second phase (**Figure 1**). The exchange between the negatively charged amino acids D75 and E75 is the most frequent among the mutations and occurs with a moderate significance (~1% rate in the second phase). Interestingly, this mutation is only specific to USA, Australia and Scotland, wherein, it occurs with the highest (0.8%) and lowest rate (0.04%) in the former and latter respectively. Strikingly, co-occurring (dependent) K141, S142 and F143 deletions which found at a less significant rate (~0.4% in the second phase) are more frequent in England, USA and Wales. More number of countries are also found to have these deletions in the second phase compared with the first phase. G82-V86 is a yet another stretch that is prone to undergo mutations. For instance, the deletion of M85 (~0.4%) is found in England, Wales, Scotland, France, Belgium, Denmark, Greece, UAE, USA, Canada and Australia with the highest occurrence in England (**Figure 1B**). Similar to K141, S142 and F143 deletions, co-existence (dependent) of G82 and H83 deletions (0.12%) is found in England, Wales, Scotland, Belgium, USA and Australia. Independent deletion of V84 and V86 is also found in certain countries (in second phase) (**Figure 1)**. Besides, V84H (England) and M85V (Wales, Scotland, Switzerland and USA) is also seen, but, with a relatively lesser rate (0.04%). The exchange between the aliphatic amino acids, V56 and I56 (with a less significance), is found to be specific for England and Wales. Other less significant mutations are, D139Y (Wales), D139N (Australia, USA, England and Wales), R124C (the highest occurrence is in England along with a single occurrence each in Russia, Ireland and India), A76T(USA), H45Y (Spain and USA), Y118C (Luxembourg and Belgium), E37K (Australia), co-existing (dependent) S135N and Y136 deletion (England, India and USA), A90V (England and Wales) and S166G (USA, Singapore, Denmark, Austria and Scotland). Notably, these significant mutations are present mainly in the loop regions of NSP1 in the I-TASSER generated structure^18^.

**Figure 1.**
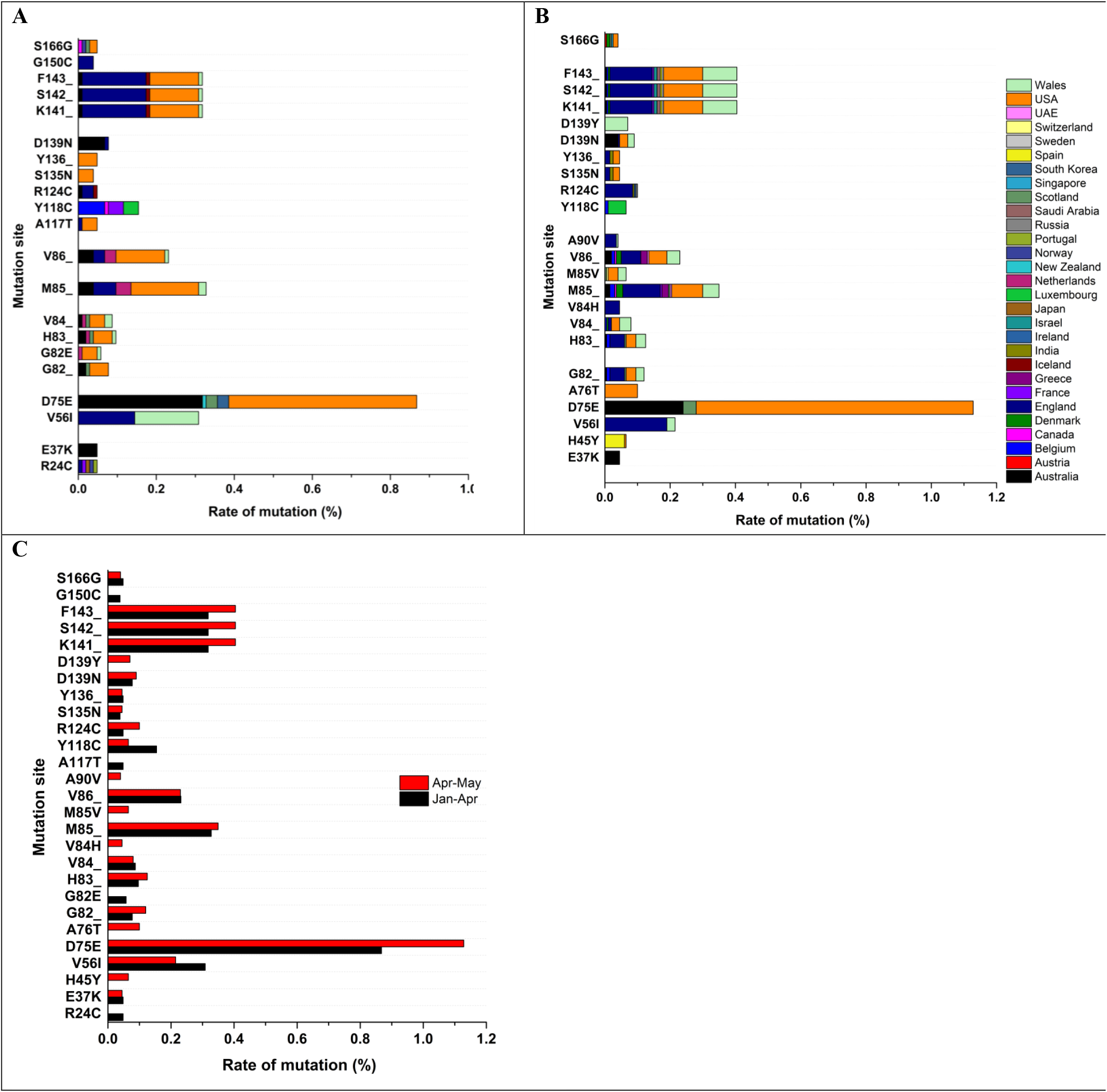
NSP1 mutation statistics. Bar diagram illustrating the demographic distribution of significant mutations found in (A) phase 1 and (B) phase 2. Note that the color coding scale represents 29 countries that have NSP1 significant mutations. In total, there are 17 mutations common to both the phases. Among them, D75E found to be a moderately significant mutation (>1%) in phase 2. C) Bar diagram showing the position-wise comparison of phase 1 (Black) and phase 2 (Red) mutation rates.

### Mutational hotspots in NSP2

NSP2 is of 638 amino acids long and interacts with the host factors and manipulates the host intracellular signaling^19^. A total of 77 positions in both the phases are prone to undergo changes (80 different changes), among which, 26 (insignificant in the phase 2) and 20 (T44I,G115C,P129L,E172K,A184S,A192V,H208Y,S263F,I296V,S369F,T388I,L400F, F437L,L462F,D464A,A476V,T547I,G548S,P568L and T634I emerge in phase 2) mutations are specific to phase 1 and phase 2 respectively. Remaining 34 mutations occur in both the phases. Hereinafter, unless otherwise mentioned, the mutation statistics are discussed only for the second phase when a mutation is present in both the phases. In phase 2, 46 countries have at least one of the 54 common significant mutations. Interestingly, T85I is the highly significant mutation in NSP2 which occurs at the rate of 22% and is significant in 37 countries in phase 2 (**Figure 2(B)**). Although Iceland, Norway, Finland and Czech Republic that do not have new data in the second phase, they do have T85I in the first phase. In total, 41 countries have this mutation indicating the global spread of I85 (54% occupancy among the second phase mutations). This mutation is found at the highest frequency in USA with nearly 56% of its sequences having this mutation. USA, Denmark, England, Australia and Israel are the top five countries possessing T85I in the second phase. Further, P585S (~3.5%), I559V (~3.3%), deletion of D268 (~2.4%) and G212D (~2.2%) are found at a moderate significant rate in the phase 2. A total of 26 countries (England, Wales, Scotland, France, Belgium, Germany, Ireland, Greece, Sweden, Singapore, Taiwan, Thailand, Vietnam, India, Jordan, USA, Australia, Chile, Iceland, Netherlands, Portugal, Finland, Canada, Ghana, Congo and Brazil) undergo P585 to S585 mutation in both the phases, among which, Iceland, Netherlands, Portugal, Finland, Canada, Ghana, Congo and Brazil have this mutation only in the first phase **(Figure 3)**. Very interestingly, the same countries undergo I559 to V559 mutation with a nearly similar rate. These two mutations occur at the highest frequency in England (~2.4-2.5%) followed by Wales (0.3%) and Australia (0.16-0.24%). They assume 8-9% and 12-13% among the phase 1 and 2 mutations respectively suggestive of their co-existence. The deletion of D268 (6% among the second phase mutants) occurs totally in 21 countries (England, Wales, Scotland, France, Iceland, Netherlands, Spain, Belgium, Finland, Ireland, Denmark, Sweden, Slovakia, Russia, Singapore, Taiwan, USA, Canada, Australia, New Zealand and Chile) together in both the phases. England is the highest (1.4%) bearer of this deletion in the second phase followed by Netherlands, Scotland and Ireland. Additionally, the exchange between G212 and D212 is found totally in 25 countries (England, Wales, Scotland, Luxembourg, Iceland, Netherlands, Portugal, Belgium, Germany, Finland, Austria, Ireland, Denmark, Greece, Hungary, Sweden, Singapore, Taiwan, Thailand, Israel, USA, Canada, Senegal, Australia and Brazil) together in both the phases. The rate of mutation of G212D is 5% among the second phase mutations. Wales has the highest (2.4%) occurrence of this mutation, followed by England and Denmark (**Figure 3**).

**Figure 2:**
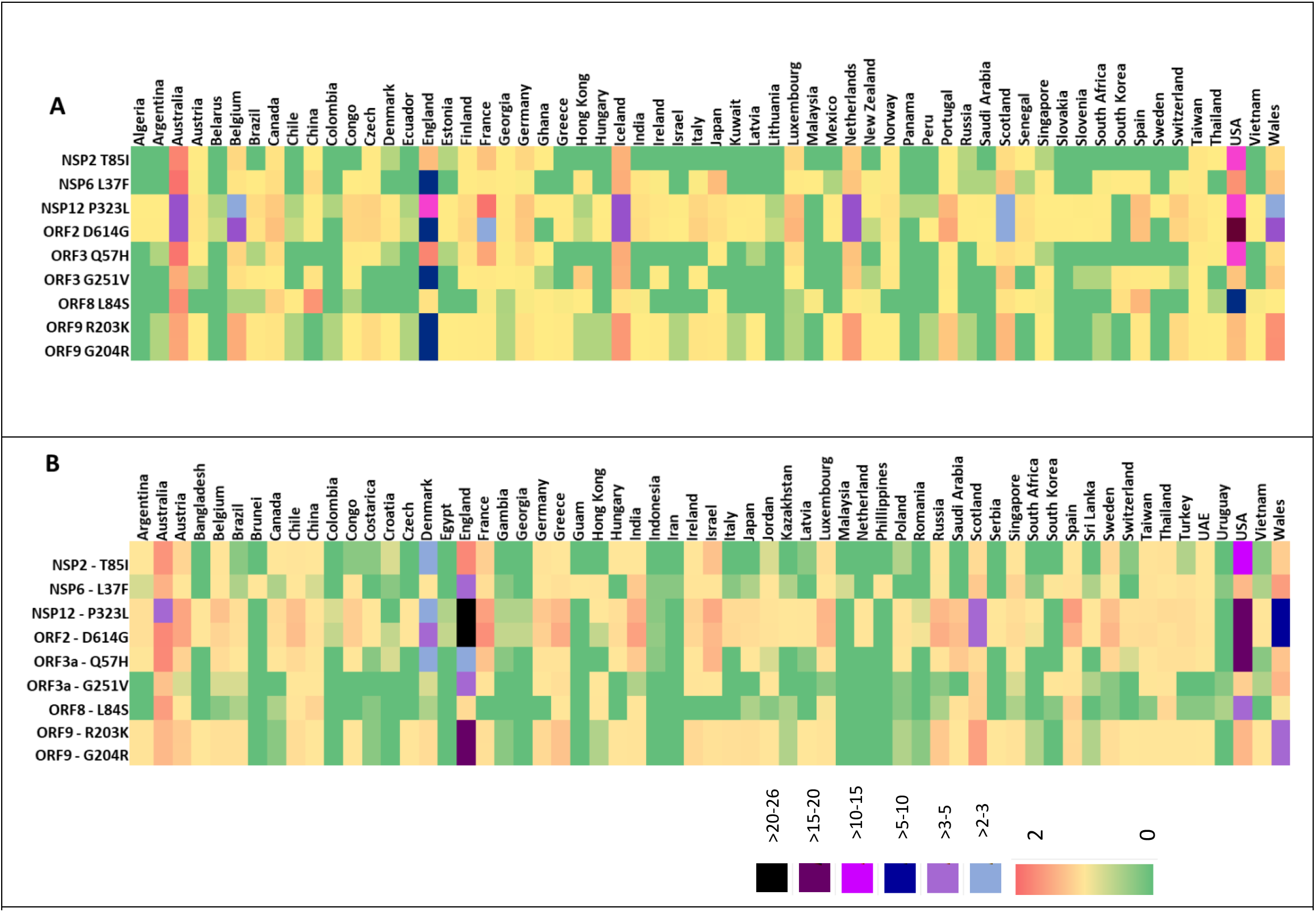
Heat map representing the country wise rate of occurrence of 9 high significant mutations in phase (A) 1 and (B) 2. A) 60 countries have at least one of the high significant mutations and totally 9 high significant mutations (>10%) have been observed in the first phase. In total, 16 countries are found to have all the mutations in phase 1. B) 61 countries have at least one of the 9 high significant mutations. Note that ORF3a-G251V and ORF8-L84S occur only at the rate of ~7% in phase 2, while the other mutations occur at a rate greater than 10%. Totally 20 countries found to have all the mutations in phase 2. Notably, namely Australia, Belgium, England, Luxembourg, Scotland, Singapore, Taiwan, USA and Wales are found to have all the mutations in both the phases.

**Figure 3:**
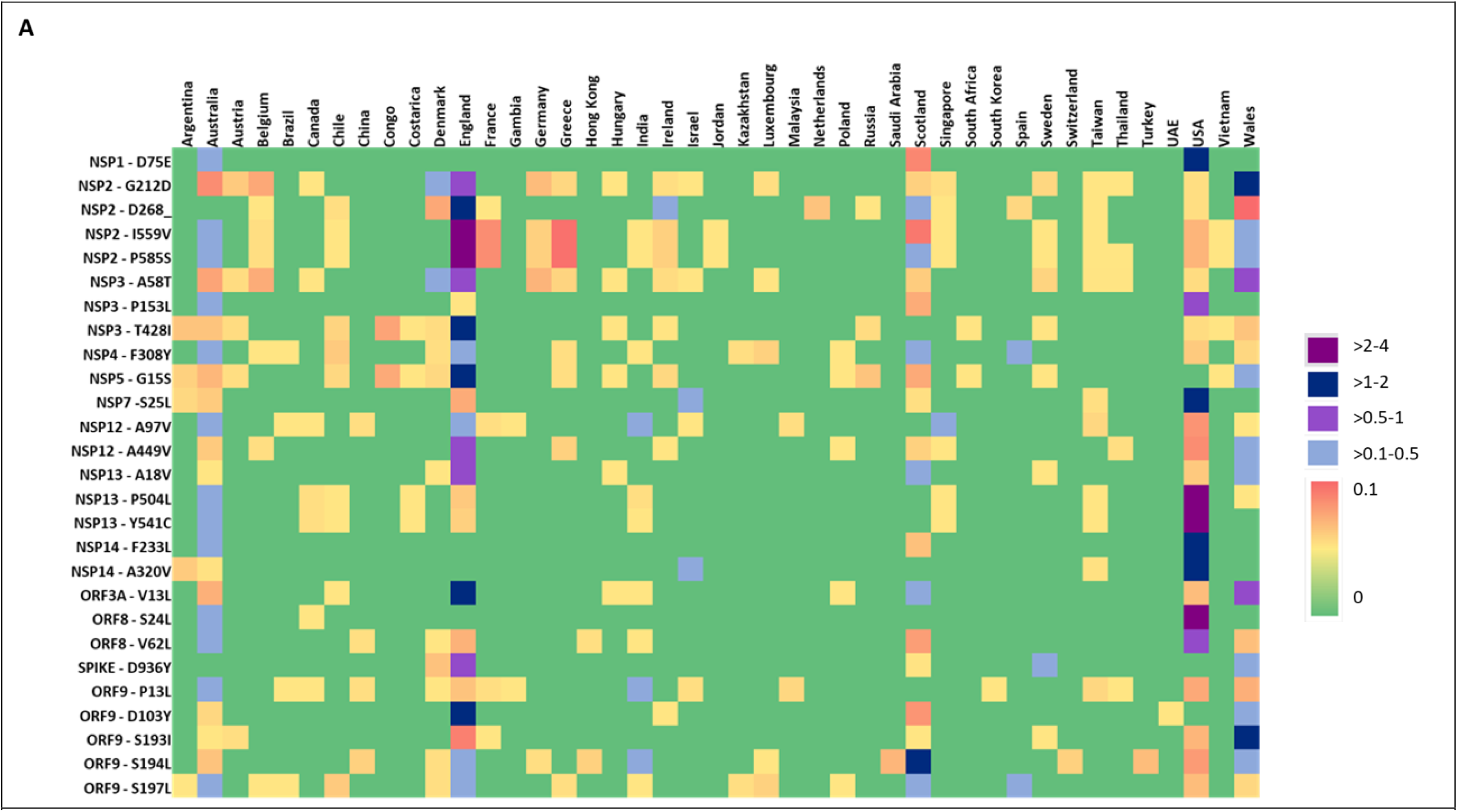
Heat map representing country wise rate of occurrence of 27 moderate significant (MS) mutations in phase 2. Note that 43 countries have at least one of the MS mutations (between 1 – 10%) in second phase. Among the 43 countries, Australia is found to have 25 out of the 27 MS mutations, followed by USA (24), England (23), Wales (20) and Scotland (20).

The remaining mutations occur at a less significant rate (below 1%). However, some of them are specific for certain countries and each assuming ~2% of the 2^nd^ phase mutations. For instance, F10 to L10 is highly specific to USA with a few incidences in Australia. Similarly, V198I has the highest occurrence in Australia along with random incidences in many countries. P91 to S91 is a yet another mutant that is found at the highest frequency in Australia followed by Scotland. T166I is highly found in Denmark with a relatively lower existence in Iceland and Latvia. H237R is seen frequently in England and Wales along with a random presence in a few other countries. H237Y is also found in England and Wales. While T371I is specific to USA, S211F has the highest incidence in England (along with a less occurrence in USA, Wales, Congo and Turkey). G339S exists with the highest frequency in England and randomly in some countries. Although the 3D structure of NSP2 is unavailable, projecting these mutations onto the I-TASSER generated structure^18^ indicates that the mutations are distributed throughout the protein.

### A58T occurs at the highest frequency in NSP3

NSP3 is a multidomain protein and is the largest (1945 amino acids) among the SARS-CoV-2 proteins. It has a ubiquitin like activity domain 1 (UBL1) (15-111)^20^, macrodomain (207-379) ^21^, ubiquitin like activity domain 2 (UBL2) (746-806) ^20^, papain-like protease (PL-pro) domain (745-1061)^22^, nucleic acid binding domain (1089-1201) and transmembrane region (1494-1563) ^22^. There are 131 positions undergo mutations at a significant rate in the second phase in 41 countries, among which, 45 mutations are common to both the phases. 49 new mutations emerge in the second phase with a low (0.05%) to moderate (0.26%) mutation rate. 43 mutations that are found in the first phase couldn’t sustain during the second phase. A58T which is present in the UBL1 domain occurs at the highest frequency in both the phases with the moderate rate of 0.7% and 1.8% respectively in the phase 1 and 2. It occurs at a rate greater than 0.6% in England and Wales, whereas found at a rate below 0.2% in the other 16 countries **(Figure 3)**. The other mutations that are found in both the phases with a mutation rate ranging between 0.3% and 1.4% in the second phase are: P153L (occurs at the rate of 0.8% and 0.2% respectively in USA and Australia), D218E (England having the highest mutation rate of 0.3%), T428I (England possessing the highest mutation rate of 1.12%), S1197R (Singapore and India having the top 2 mutation rate of 0.3% and 0.08% respectively), T1198K (India and Australia having the top 2 mutation rate of 0.4% and 0.2% respectively), P1326L (Ireland having the highest mutation rate of 0.2% and P1326T (0.07% is also found in certain countries)), S1424F (Wales having the highest mutation rate of 0.4%) and A1431V (the highest rate of 0.12% is found in Denmark). Notably, these mutations do not occur in the PL-pro domain (PDB ID: 6WUU), which makes this domain to be a potential drug target in line with the earlier attempts ^2^ (**Figure 4)**.

**Figure 4:**
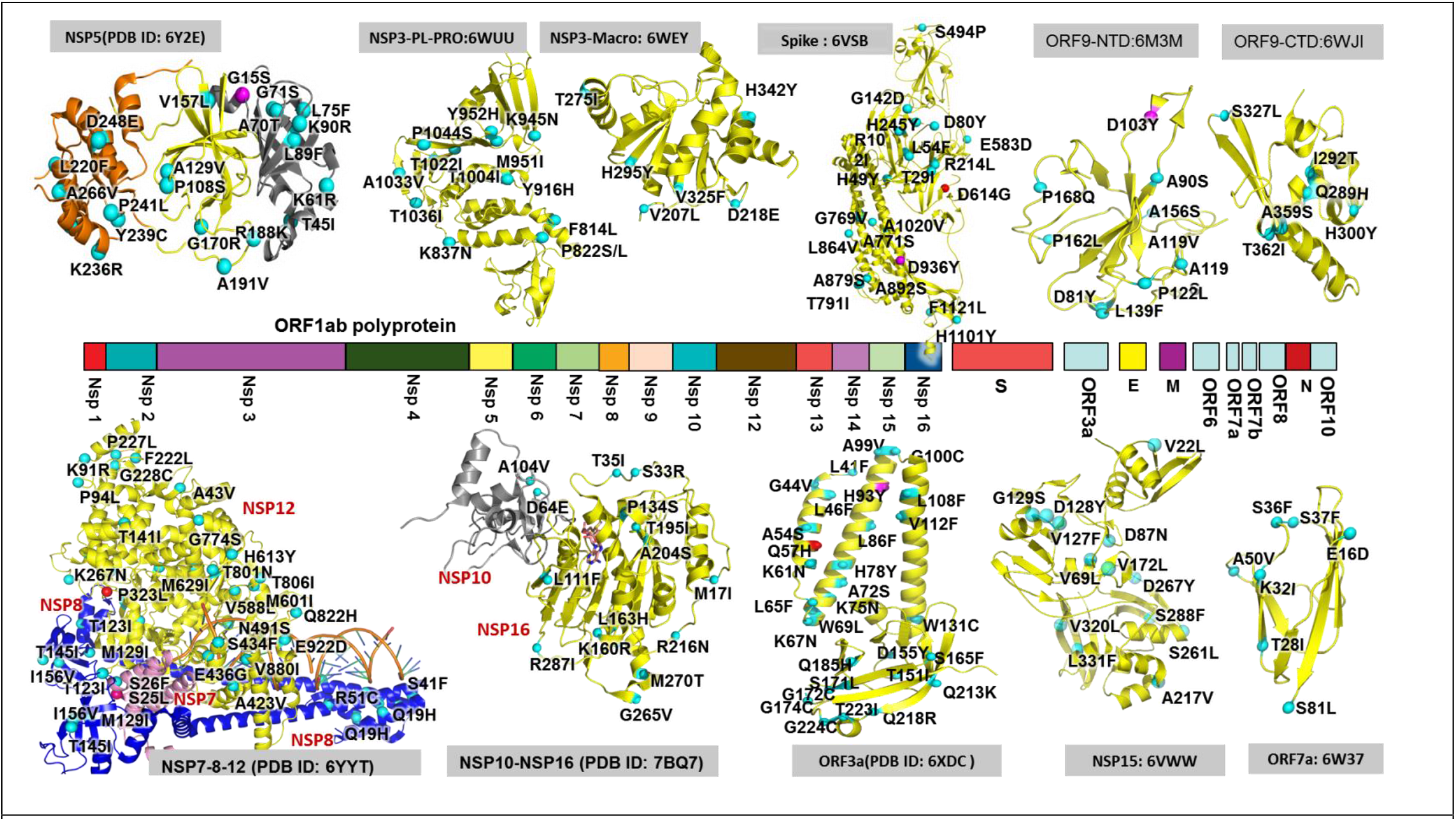
Projection of SARS-CoV2 mutations onto the experimentally known proteins structures. The high (HS), moderate (MS) and Low (LS) significant mutations are colored red, magenta and cyan respectively. Note that the cartoon representation of the protein and corresponding PDB IDs are indicated adjacent (top and bottom) to the corresponding coding region (middle). Note that NSP3-PL-PRO and NSP3-Macro indicate PL-PRO and macro domains of NSP3 respectively. Similarly, ORF9-NTD and ORF9-CTD indicate N-terminal and C-terminal domain of ORF9. For the sake of clarity, only the monomeric structures of NSP3, spike, ORF3a and ORF9 are given.

Apart from these, K19R (specific to England), T73I (the highest rate of 0.12% in England), S166G (specific to England, Scotland and Wales), T181I (present in 5 countries and T181A is specific to Wales), P395L (occurs in 4 countries), G489D (exclusive to England), L557F (found primarily in Thailand), I707V (the highest in USA), Y916H (specific to Scotland and Wales), Y952H (specific to Wales), S1285F (the highest in England followed by Scotland and 5 other countries), T1303I (specific to England) and M1436V (found largely in England) occur with a low significant rate (between 0.1% and 0.3%).

### NSP4 exhibits less number of mutations

NSP4 is 500 amino acid long protein, having multiple transmembrane domains, involved in double membrane vesicles (DMVs) formation for compartmentalization of virus replication with the help of NSP3 and NSP6^23^. Among the 16 mutations that occur in the first phase, 10 of them are common to both the phases (M33I, L206F, T295I, A307V, F308Y, G309C, H313Y, L360F, L438F and A457V). Additionally, V20F, M33L, S34F, L111S, T214I, T237I, L264F and S481L emerge in the second phase. The exchange between the aromatic amino acids F308 and Y308 is the highest among the mutations and it occurs at rate of ~2% in the second phase. Australia has the highest occurrence of F308Y followed by Scotland, England and Spain. Wales, Luxembourg, Belgium, Denmark, Greece, Poland, India, Kazakhstan, USA, Brazil and Chile also have this mutation in the phase 2 **(Figure 3)**. The remaining common mutations occur below 1% mutation rate and hence are not discussed. Among the common mutations, T295I, A307V, F308Y, G309C and H313Y occur in the helix-turn-helix motif (in the I-TASSER generated structure ^18^), wherein, T295I and H313Y exist in the middle of 2 different alpha helices and the others occur in the turn of the motif. M33I/L is present in the loop region that follows the N-terminal alpha helix.

### Conservation of NSP5 catalytic site and N-terminal finger

NSP5 also known as main protease (M^pro^) or 3 chymotrypsin-like protease (3CL^pro^) plays a major role in cleaving the virion polypeptide chain 1a and/or 1ab to produce the functional forms of NSP4 to NSP16^24^ which are crucial for the viral replication in the host. Thus, it is an attractive target to treat SARS-CoV-2. It is a 306 long amino acid protein and forms a homodimer for the proteolytic activity function and each subunit contains a His 41/Cys145 catalytic dyad. It shares 96% sequence identity with SARS-CoV. NSP5 has three different domains namely, the chymotrypsin-like (domain I) and picornavirus 3C protease–like (domain II) domains (residues 10 to 99 and 100 to 182, respectively) and domain III (residues 198 to 303), which are involved in regulating the dimerization of the main-protease^24^. There are 10 different positions undergo significant changes in both the phases (G15S, K61R, G71S, L89F, K90R, P108S, A191V, L220F, K236R and A266V). New mutations that evolve as significant mutations in the second phase are T45I, A70T, L75F, P108L, A129V, V157L, G170R, R188K, Y239C, P241L, D248E and N274D. Together in both the phases, 32 countries have at least one of the significant mutations in NSP5 (Luxembourg, Iceland, Netherlands, Germany, Finland, Georgia, Brazil, England, Wales, Scotland, France, Spain, Austria, Ireland, Denmark, Greece, Hungary, Sweden, Poland, China, Japan, Russia, Taiwan, Vietnam, India, Jordan, USA, Costa Rica, South Africa, Congo, Australia, Argentina and Chile).

Among these, G15S and K90R located in the chymotrypsin-like domain are occurring at a moderate significance (1-2% mutation rate). G15S appears 38% among the second phase mutations with England and Wales taking up the top 2 positions along with a few more countries in both the phases **(Figure 3)**. Iceland is the highest bearer of K90R in phase 1 (0.9% mutation rate), for which the data is not available in the phase 2. However, England and USA have significant occurrence of K90R in phase 2. Further, L89F (mutation rate of 0.5%) is found to be specific for USA. Other mutations that occupy greater than 2% among the second phase mutations are: T45I (USA and Taiwan), A70T (England and Wales), G71S (Chili, USA, Denmark, Australia and England), L75F (USA and India), P108S (predominantly in Wales and USA, and P108L is also seen predominantly in USA), L220F (USA and Scotland), D248E (the highest in Scotland), A266V (USA, Australia and Jordan) and N274D (the highest in Scotland). Luckily, the structure of M^pro^ is available (**PDB ID:** 6LU7) which indicates that these mutations do not occur either at the catalytic site or at the N-terminal finger (important for the recognition of the target protein) or at the dimeric interface indicating the conservation of its functional mechanism (**Figure 4**). These mutations in general occur on the surface exposed regions of NSP5, thus, may have an influence on the interaction with other virion protein(s) as well as the host protein(s).

### L37F is the dominant mutation in NSP6

NSP6 is a trans-membrane protein of 290 amino acid length and is presumed to participate in inducing double membrane vesicles together with NSP3 and NSP4^25^. There are 8 mutations (T10I, insertion of a F in the F34-F36 stretch, L37F, A46V, L75F, V149F, L260F and G277S) prevail in both the phases. Some mutations (K109N (England), V145I (England), Y153C (China), N156D (Netherlands and England), L167F (Australia and France), A186T (England) and G188V (Australia)) that occur in phase 1 do not present in phase 2 or vice-a-versa (L22F, V84F, A136V, M183I, G188S and I189V). L37F occurs with the highest significance (greater than 80% among NSP6 mutations in both the phases and with a mutation rate >10%) (**Figure 2(A))**. A total of 67 countries in both the phases possess these mutations. However, the other mutations occur at a less significant rate (below 1%) and some of them are specific to certain countries. Interestingly, insertion of a F amidst 3 F’s (F34,F35 and F36) is found in England, Wales, Australia, Congo, Jordan, Philippines, Qatar, Taiwan and UAE in second phase. This mutation is also found in Switzerland in the first phase. G277S is specific to Wales and L75F is found predominantly in England and USA. Similarly, T10I is specific to England, Scotland and Ireland and A46V is predominantly seen in Australia. Further, L260F is found highly in England and V149F is dominant in USA. It is noteworthy that the majority of the amino acids mutates either to F or to a more hydrophobic amino acid implicating the role of this mutations in host membrane manipulations.

### Mutations in the replication machinery

NSP7 to NSP16 are the replicase polyproteins and crucial for the protein replication. NSP7 along with NSP8 are involved in stimulating the polymerase activity of NSP12 ^26^ and is of 83 amino acid length. There are only 2 significant mutations in both the phases, namely, S25L and S26F (**Figure 3**). While the former occurs at moderate significant rate (1.5%), the latter occurs at a low significant rate (0.25%). Interestingly, USA is the highest bearer of both the mutations. Israel also has significant population of the former. These mutations occur at the turn region of the alpha helical bundle of NSP7 which interacts with NSP8 (**Figure 4**).

NSP8 forms heterodimer complex with NSP7 and acts as a cofactor for the function of NSP12^26,27^. The mutations at M129I (0.5%) and R51C/L (~0.1%) are found in both the phases, but, with a less significance. While M129I is specific to USA (along with an occurrence in Australia), R51C is specific to England and Wales). R51L is also found in England, Wales and USA in the second phase. NSP8 is a 198 amino acid long protein and rich in alpha helical content. Notably, R51 plays a major role in interaction with the double stranded RNA at the minor groove side and M129 occurs at the interface of NSP8-NSP12 (**Figure 4**, **PDB ID: 6YYT**)^28^ indicating that these mutations may have an influence on the viral replication mechanism. The other mutations that prevail either in the first (A16V, A21V, T89I, S177L and T187I) or in the second phase (Q19H, V34F, S41F, T123I, T145I and I156V) have a low significant rate (less than 0.1%).

NSP9 is a single-stranded RNA binding protein^29^ and is of 113 amino acids long. There is no common mutation found between both the phases. Further, the mutations that found only in the first phase and not in the second (T24I and P71S) or vice-a-versa (T19K, L42F, T62I, P71Q, and M101V) exhibit only a low mutation rate (between 0.04% and 0.06%).

NSP10 functions as a stimulator of NSP14^30^ and also involved in activating NSP16^31,32^. It is a 139 long amino acid protein and the rate of mutation is low as the NSP9. The exchange between acidic D and E at the 64^th^ position (England, Wales, Scotland and Ireland) occurs at the highest rate (0.2% in the second phase) and is the only NSP10 mutation found in both the phases. The other second phase mutations (A104V (England, USA, Australia) and P136S (England, Austria)) as well as A32V (first phase) are found at a low significance (0.08%) (**Figure 4**, **PDB ID: 6ZCT, 7BQ7**).

Intriguingly, NSP11 is 13 amino acids long poly-peptide and doesn’t exhibit any significant mutation.

NSP12 is the RNA dependent RNA polymerase (RdRp), the essential component of the replicase machinery. It is 932 amino acids long protein. It has 17 mutations in common to both the phases (T26I, D63Y, A97V, T141I, P227L, P323L, M380I, A423V, S434F, E436G, A449V, N491S, H613Y, S647I, M666I, G774S and T806I). Alarmingly, P323L that is located in the proximity of NSP8 binding site (**Figure 4**, **PDB ID: 6YYT**) is found in 71 countries (except Brunei, Croatia, Iran, Malaysia, South Korea, Nepal, Philippines, Turkey, Pakistan, Tunisia, Guam and Uruguay) out of 84 countries considered here with a high significance (75% mutation rate in the second phase) **(Figure 2)**. Prevalence of this mutation is also reported earlier^7,33–35^. Next, A97V (England, India, Singapore and Australia taking the top 4 positions among the 14 countries) occurs at a moderate significance (1.6%) in the second phase **(Figure 3)**. Further, T141I (the highest in Scotland followed by England along with random incidences in several countries), S434F (0.4%, the highest in Wales followed by England and a single incidence in Sweden), A449V (the highest in England followed by Wales along with some random incidences in a few countries) and M666I (0.3%, prevalent in Wales, Scotland and England alongside with a few incidences in 3 countries) are seen with a low significant rate (between 0.3 and 1%). The other common mutations are found at rate less than 0.3%. There are mutations that occur in the first phase (P21S, G25Y, G44V, T85I, M124I, T248I, R249W and R279S) become insignificant in the second phase (below 0.04%). Similarly, some mutations (S6L, A43V, K91R, P94L, I171V, F222L, G228C, T252N, K267N, L329I, V588L, M601I, T801N, T806I, Q822H, V880I and E922D) emerge in the second phase (below 0.04%).

The helicase NSP13^36^ is of 601 amino acids long and comprises of zinc binding domain (residue numbers 1-99), stalk (100-149), 1B (150-260), 1A (261-441) and 2A (442-596)^37,38^ domains. It exhibits 36 significant mutations in the phase 2, among which, 20 of them are common to both the phases (A18V, V49I, S74L, P78S, D204Y, G206C, V226L, H290Y, L297P, V348L, P364S, R392C, S468L, V479F, S485L, P504L, Y541C, V570L, M576I and T588I). The co-mutation P504L (nucleic acid binding channel, 2A domain) and Y541C (nucleic acid binding pocket, 2A domain) ^37,38^ occurs not only at the highest significance, but also, at a similar frequency in same countries **(Figure 3)**. The USA is the highest bearer of this mutation followed by Australia along with a few incidences in 7 other countries in phase 2. The exchange between the aliphatic A18 and V18 is found at the moderate significance (1.1%) and England sees the highest exchange rate (0.7%) followed by Wales and Scotland **(Figure 3)**. Followed by this, R392C possesses the mutation rate of 0.6% in the first phase and is mostly found in Netherlands sequences. However, due to a less number of data for Netherlands in the second phase, it is found at a low significant rate. Similarly, L176F that is specific to the Netherlands (0.2%) is absent in the second phase. A few occurrences are also found in other countries. Besides, L297P (0.2%) is found at the highest rate in Scotland and Spain along with England, Wales and Luxembourg. Besides, P47S, P53S, T127I, S166L, V169F, P172S, V193I, A237T, Y253H, D260Y, A338V, Y396C, L428F, T481M, K524N and A598V are the significant mutations that evolved in the 2^nd^ phase. Strikingly, the USA possesses the majority of these mutations by having 31 of the 2^nd^ phase mutations (*viz*., except A237T, V348L, S485L, V570L and T588I).

NSP14 is an exonuclease with proofreading activity ^30,39^. It is a 527 long amino acid protein that has 7 significant mutations common between the 2 phases: M72I, A225V, F233L, T250I, A320V, P443S and A482V. Among these, F233L and A320V are seen at a moderate significance (~1.4%) **(Figure 3)** and are found in the exoribonuclease (ExoN) and (guanine-N7) methyl transferase (N7-Mtase) domains of NSP14 respectively ^30^ (modeled with SWISS-MODEL based on SARS-CoV NSP14 structure with sequence identity of 95% (PDB ID: 5C8S). In the phase 2, F233L is highly found in USA followed by Australia and Scotland. A320V is highly found in USA followed by Israel along with a few incidences in Argentina, Australia and Taiwan. Next, T250I (found in Australia, England, Scotland, Spain, Sweden and USA) and A225V (found in England and Wales) are found at 0.3% mutation rate. The remaining common mutations along with the newly emerged second phase mutations, H26Y, V182L, T193A, S255I and M501I have a low significant rate (~0.05%). T16I, P46L, P121L, I150T, L177F, P203L, R205C, T206I, P297S, P327S and L493F are found in the first phase but not in the second phase.

The uridylate-specific endoribonuclease^40^ protein (346 amino acids, (NSP15)) has 8 mutations common to both the phases: V22L (the highest occurrence in Australia, along with USA, England, Scotland and Wales), V127F (the highest occurrence in USA with a single occurrence each in England and Wales), G129S (specific to USA), A217V (England, Wales, Scotland, Ireland and Singapore), D267Y (Australia, Denmark, Sweden, India in the second phase) and V320L (the highest occurrence in England along with Wales, Scotland, USA, Australia and Saudi Arabia). While V320L, A217V and V22L have a mutation rate in the range of 0.3% and 0.8%, the remaining mutations have a rate of mutation below 0.1%. V172L (the highest incidence in Thailand along with USA, Japan, England and Wales) emerges in the second phase with mutation rate of 0.3%. Interestingly, D219N that occurs at a rate of 0.4% in the phase 1 (the highest in Netherlands followed by Iceland) is insignificant in the phase 2 either due to the unavailability of sequences (Iceland) or due to lack of sufficient data (Netherlands).

NSP16 is 298 amino acids long ribose-2’-*O*-methyltransferase^41^. All the mutations in both the phases have only low (0.04%) to moderate (0.35%) significance. There are 7 significant mutations that occur in both the phases, *viz*., S33R, T35I, P134S, K160R, L163H, T195I and G265V. While the rate of mutation for S33R (the highest in the USA along with Israel, Australia, Russia and England), P134S (the highest in the England along with Australia, Vietnam and Chile) and K160R (the highest in the England along with USA, Wales, Belgium, Hungry, Sweden, China, Saudi Arabia, Singapore, India and Australia) is found to be ~0.3% in the second phase, the others (T195I (specific to Wales and England), G265V (specific to USA and turkey), T35I (highest in England along with some other countries) and L163H (specific to USA)) show mutation rate between 0.05% to 0.1%. There are some mutations (P12S, Q28K, G77R, G113C, G165C, K182N and V294F) that occur only in the phase 1 (found in the range 0.04% and 0.1%). Interestingly, P134S is found in the cavity that is in proximity to the SAM binding site of NSP16 **(Figure 4, PDB ID: 7BQ7)**.

### Proteins coded by subgenomic mRNA of SARS-CoV-2

The subgenomic mRNAs encodes the structural proteins, namely, spike (S) (ORF2), envelope (E) (ORF4), membrane (M) (ORF5), nucleocapsid (N) (ORF9/nucleoprotein) and the accessory proteins, namely, ORF3a, ORF6, ORF7a, ORF7b, ORF8 and ORF10.

### D614G and D936Y are the prominent mutations in the spike protein

The spike protein is a glycoprotein that spans the viral membrane and the extracellular region. It mediates the viral entry by interacting with the host-cell receptor, namely, angiotensin-converting enzyme 2 in the same way as the other betacoronaviruses ^42–46^. It is encoded by ORF2 of the viral genome and is one of the four structural proteins present in SARS-CoV-2. It is of 1273 amino acids long and exists as a trimer^47^. It has two subunits, namely, S1 (amino acid residues 1 to 685) and S2 (686 to 1273). The subunit S1 consists of a signal sequence (1-12), an N-terminal domain (13-305) and a receptor binding domain (330-521). The subunit S2 has a fusion peptide (788 to 806), a heptad repeat 1 (912-984), a heptad repeat 2 (1163-1213), a transmembrane domain (1213-1237) and a cytoplasmic tail (1238-1273)^47^. As reported in the earlier studies ^48–51^ D614G (present in the S1 subunit flanked by the receptor binding site) **(Figure 4)** mutation is found to be highly significant and present at 75% rate in the second phase. Together in both the phases, 68 countries (except 16 countries, namely, Brunei, Sweden, Panama, Cambodia, Tunisia, Malaysia, Nepal, South Korea, Philippines, Iran, Pakistan, Ghana, Uruguay, Qatar, Bangladesh and Guam) have this mutation **(Figure 2(B))**. Followed by this, D936Y (heptad repeat 1) assumes a moderately significant rate (1.2% in the second phase) with England, Wales and Sweden taking the top 3 positions along with 2 other countries **(Figure 3)**. Amongst the rest of the 22 mutations that are common to both the phases, L5F (domain S1, occurs in US, England and Wales with a rate of at least 0.1% each), R21I (domain S1, Wales and England have the mutation rate of 0.5% and 0.2% respectively) and P1263L (cytoplasmic tail, the highest in England and Wales with 0.4% and 0.1% rate respectively) exhibit mutation rate greater than 0.6% in the phase 2. The other common mutations having rate between 0.1% and 0.6% in the 2^nd^ phase are found in the S1 domain: L18F (England having 0.1%), H49Y (China having 0.09%), L54F (England having 0.2%) and H146Y (USA having 0.3%). 6 of the newly emerged mutations in the second phase also exhibit mutation rate between 0.1% and 0.2%: R102I (Wales having the highest rate of mutation), R214L (England having the highest rate of mutation), T478I (specific to England), A829T (specific to Thailand), R847K (specific to Wales) and A1020V (random occurrences). 19 new mutations in the second phase have a mutation rate lesser than 0.1%. A831V (the highest occurrence in Iceland), S943P (specific to Belgium) and G1124V (specific to Australia) seen in the first phase with a mutation rate of ~0.2% are not found in the second phase. Similarly, 27 mutations that have a mutation rate lesser than 0.2% in the first phase are not seen in the second phase. Notably, an insertion of Y in between 264A and 265Y (S1 domain) is found to be specific for England in both the phases (0.01%).

### Q57H and G251V are prominent in ORF3a

ORF3a is an accessory protein of 275 amino acids long. It is a transmembrane protein which can exist in dimeric or tetrameric forms, and are shown to have ion channel activity^52–54^. While there are 23 new significant mutations emerged in the second phase, 12 mutations that are significant in the first phase are not seen in the second phase. 21 significant mutations are common to both the phases. V13L, Q57H and G251V occur at a high significant rate in the phase 2 which are happened to be the common mutations. Q57H occurs at the highest rate of 27% in phase 2 and a total of 50 countries in both the phases have this mutation. Among these countries, USA has the highest mutation rate of 15.3% in the second phase followed by England, Denmark and Australia (each greater than 1%) **(Figure 2(B))**. G251V occurs at the rate of 12% in 30 countries in the second phase with England taking the top position in the mutation rate (4.3%) followed by Wales, Australia, USA and Scotland. V13L occurs at the highest rate in England in the second phase followed by Wales and Scotland in addition to a few incidences in 6 more countries **(Figure 2(B))**. T14I (Scotland and Canada have the ~0.1% mutation rate along with a few occurrences in the England, Luxembourg, USA and Belgium), L46F (the highest in England with the 0.2% rate), A54S (the highest in Wales with 0.6% rate and 0.1% in England), H93Y (0.5% in Wales), A99V (the highest rate of 0.2% in England along with 7 more countries), T175I (England and USA having the mutation rate of ~0.1%), Q185H (the highest mutation rate of 0.2% in Wales followed by 3 other countries) and G196V (England and Spain having mutation rate of ~ 0.1%) are the other mutations that occur at a rate ranging between 0.1% and 1% in the second phase. The remaining mutations in the phase 2 occur at a low significant rate (0.05%) and are not discussed here.

### The envelope protein undergoes mutation at a low rate

The envelope protein, which is encoded by ORF4, is of 75 amino acids long and functions as viroporin and has a vital role in viral assembly^55^. Intriguingly, the rate and the number of mutations are very less in the envelope protein. For instance, there are only five mutations in the second phase S68F, P71L, D72H, L73F and V75L each occurring at a low mutation rate (0.04%-0.1%). Each of these mutations occurs randomly in at least 4 countries. Notably, D72H and L73F are found in both the phases.

### T175 is a moderately significant mutation in the M-protein

ORF5 encodes for the membrane protein that is involved in viral assembly^56^ and is a 222 amino acids long protein. Although D3G, V70I, W75L, T175M and S214I are found to be common to both the phases, T175M occurs at a moderate significance rate (~3% in 1^st^ phase and ~1% in second phase). It occurs at rate a greater than 0.1% in England and Austria followed by 14 other countries in second phase. Additionally, Iceland and Netherlands also have a mutation rate greater than 0.7% in the first phase whose data are either not available (Iceland) or significantly less (Netherlands) in the second phase. D3G occurs at a low significance (0.7%). It occurs at the highest rate in England (0.4%), while the remaining 18 countries bear 0.3% in second phase. The remaining 3 common mutations, 4 mutations found only in the first phase and 13 newly emerged mutations in the second phase have a mutation rate of less than 0.1% each.

### ORF6 protein undergoes mutations at a low significance

ORF6 encodes one of the accessory proteins (61 amino acids long) of SARS-CoV-2 and is found to be the most potent interferon antagonist^57,58^. Like envelope protein, it has a very low mutation rate. W27L, I33T and K42N are the common significant mutations in both the phases. W27L and I33T occur at a higher rate (~0.15%) compared with others (lesser than 0.05%) in the phase 2. W27L occurs at the highest rate in England along with Wales, Scotland, Australia and Singapore. I33T is found in 7 countries in the second phase. R20S that is found as a significant mutation in Senegal in the first phase is not seen in the second phase as there is no data available for Senegal in the second phase. However, D53G and D53Y emerge as new mutations in the second phase with mutation rate of 0.04% each.

### Proteins encoded by ORF7a and ORF7b have mutations of low significance

ORF7a encodes for 121 amino acids long protein and is one of the virulence factors involved in interfering with host’s innate immunity^59,60^. Among the six significant common mutations (T28I, A50V, S81L, V93F, L96F and V104F) found in the ORF7a, S81L has the highest mutation rate of 0.4% in phase 2. S81L is found in 3 countries in the following order of mutational rate: Australia > USA > Scotland. The other common mutations occur at a rate lower than 0.1% in the second phase. A8T, E16D, K32I, S36I, S37F, V108L and T120I emerge in the second phase, but, with a mutation rate lesser than 0.07%.

Interestingly, ORF7b protein (43 amino acids long) doesn’t have any significant mutations during the first phase. However, L20F, S31L, T40I and C41F are found to be significant in the second phase, but, with a lower mutation rate (below 0.25%). Among these, C41F has the highest mutation rate of 0.2% with Thailand having 0.1%.

### L84S is a high significant mutation in ORF8

S24L, A51V, V62L, A65S, S67F, L84S, F120L and a deletion in ORF8 are found to be common to both the phases. The protein encoded by ORF8 has 121 amino acids and presumed to be responsible for host immune evasion^61,62^. While L84S occurs at a high significance (14% in phase 1 and 8% in phase 2), S24L (2% in phase 2) and V62L (1% in phase 2) occur at a moderate significance **(Figure 2, 3)**. A total of 44 countries together in both the phases bear L84S mutation with USA having the highest mutation rate (5%) followed by Australia (1%) in the second phase. Further, England, Scotland, Spain, China, Saudi Arabia and Thailand have overall mutation rate above 0.1% in the second phase. USA is the highest bearer of S24L followed by Australia and Canada in both the phases. Similarly, USA has the highest rate of V62L followed by Australia along with a few incidences in some countries. Notably, a deletion in ORF8 is found in Singapore and Taiwan in both the phases, encoding only 9 amino acids long peptide. Remaining 4 common mutations exhibit only a low mutation rate (lesser than 0.25%) in both the phases. Similarly, 4 new mutations in the second phase have a low significance (lesser than 0.1%). Further, 7 mutations which are present in the first phase do not occur in the second phase.

### The co-mutations R203K and G204R occur in nucleocapsid protein with a high significance

ORF9 of SARS-CoV-2 encodes for nucleocapsid (N) protein of 419 amino acids long, which is a multidomain and multifunctional protein and is vital for viral genome packaging. N protein comprises of three domains: N-terminal RNA-binding domain (NTD) (49-174), an intrinsically disordered central Ser/Arg (SR)-rich linker(175-246) and C-terminal dimerization domain (CTD)(247-365), flanked by the disordered regions at both N and C termini^63–65^. There are 36 significant mutations found to be common between the 2 phases (P13L, D22G, T24N, S33I, D81Y, D103Y, A119S, A119V, P122L, L139F, A156S, S180I, S183Y, R185C, S188L, S188P, S193I, S190I, S194L, R195I, S197L, P199S, S202N, R203K, R203S, G204R, T205I, A208G, M234I, G238C, Q289H, I292T, S327L, K373N, D377Y and T393I) across 68 countries. Among these, R203K and G204R are not only found at a high significance (~27%) but also act as co-mutations (dependent). Both of them co-exist in 48 countries and occur at a high mutation rate of 15.4% in England followed by Wales (4.3%) and Scotland (1.1%) in the second phase. The remaining countries (France, Luxembourg, Spain, Belgium, Germany, Italy, Czech Republic, Austria, Ireland, Denmark, Greece, Hungary, Sweden, Poland, Turkey, Switzerland, Serbia, Romania, China, Japan, Russia, Singapore, Taiwan, Thailand, Vietnam, India, Israel, UAE, Jordan, Hong Kong, Bangladesh, Kazakhstan, Sri Lanka, USA, Canada, Costa Rica, South Africa, Congo, Gambia, Australia, Argentina, Brazil Chile, Ecuador and Estonia) in the second phase possess ~6% of this mutation **(Figure 2)**. Interestingly, R203S that is found at a low mutation rate (0.06%) doesn’t accompany with the mutation at G204. Notably, S180, S183, S187, S188, S190, S193, S194, S197 and S202 present in a serine rich stretch (SR rich linker domain, residue number 180-202) change primary to aliphatic/aromatic amino acids I,Y,L,L/P,I,I,L,L and N respectively. Among these, S193I, S194L and S197L have mutation rates between 1-2% (moderate significance) while S188L and S190I are found at the rate of 0.6% and 0.3% respectively. Indeed, certain countries have more than 50% of these mutations: S188L (the highest in Scotland followed by England), S190I (the highest in England), S193I (the highest in Wales followed by England), S194I (the highest in Scotland followed by England, India and Wales) and S197L (the highest Australia followed by Scotland, England and Spain). Apart from these, P13L (N-terminal disordered region, the highest in Australia), D22G (the highest in England followed by Australia and Wales) and D81Y (the highest in England) also occur at a moderate significant rate (1%). T205I (the highest in England), A208G (the highest in USA along with an incidence in Australia), I292T (the highest in Brazil) and T393I (the highest in the USA along with 1 and 2 incidences in Australia and England respectively) occur at a rate ranging from 0.2 to 1%. 20 new mutations emerge in the second phase, out of which, D401Y, R209I, R209T, T247I, K374N and T391I occur at a rate ranging between 0.05% and 0.25%. Notably, 18 of the 36 significant mutations (including R203K and G204R co-mutations) in the second phase occur in the stretch S180-R209 of SR rich linker region, for which, the structural information is unknown **(Figure 4)**.

### The protein encoded by ORF10 undergoes changes at an insignificant rate

The protein encoded by ORF10 is of 38 amino acids long. While the first phase has 4 significant mutations, the second phase has 6 significant mutations. But, the rate of mutation is less than 0.15% in both the phases. Among these, P10S, S23F and R24C are common to both the phases. These mutations are randomly seen in anyone of the following countries: England, Wales, Scotland, Spain, Denmark, Israel, USA and Australia.

## Discussion

The SARS-COV-2 pandemic is a major threat to the public health. Several attempts are being made to obtain a potential drug molecule or vaccine candidate with a broad coverage. To this end, a comparative analysis of SARS-CoV-2 whole genome protein sequences has been performed here by considering 31389 whole genome sequences from a diverse clinical and geographical (84 countries) origin to identify the mutations that occur in 26 SARS-CoV-2 proteins. Prior to the analyses a manually curated database has been created for the whole genome protein sequences and the analyses have been done by dividing the dataset into 2 different phases (phase 1: Jan-April 17,2020 and phase 2: April 18-May17, 2020).

The results indicate that a total of 563 (second phase) and 534 (first phase) significant mutations are found across the SARS-CoV-2 proteins in 84 nations (**Figure 6**), among which, 9 (occurring above 10% in anyone of the phases), 27 (occurring between 1 and 10% in second phase excluding G251V-ORF3a and L84S-ORF8 as they occur greater than 10% in phase 1) and 527 (occurring between 0.4% and 1% in the second phase) are high, moderate and low significant mutations respectively (**Figure 6**). The high significant mutations include T85I (~22% across 37 countries in phase 2, NSP2, ORF1ab), L37F (10% across 47 countries in phase 2, NSP6, ORF1ab), P323L (~75% across 53 countries in phase 2, NSP12, ORF1ab), D614G (~75% across 54 countries in phase 2, spike), Q57H (~27% across 41 countries in phase 2, ORF3a), G251V (~7% across 31 countries in phase 2 and ~13% across 33 countries in phase 1, ORF3a), L84S (~8% across 32 countries in phase 2 and ~13% across 32 countries in phase 1, ORF8), R203K (~27% across 48 countries in phase 2, nucleoprotein) and G204 (~27% across 48 countries in phase 2, nucleoprotein) (**Figure 2**). Interestingly, R203K and G204R are co-occurring (depending) mutations as they occur in nearly similar frequency in same countries. Among all the highly significant mutations P323L-NSP12 and D614G-spike are found in majority of the countries. Intriguingly, some of these mutations are highly prevalent in certain countries compared to rest of the mutations: P323L, D614G, R203K and G204R in England, P323L, D614G and Q57H in USA. The occurrence of the above mutations in both the phases with a high significant rate clearly indicates that these mutations might have developed very early and may have a positive selection pressure. The presence of all these mutations in China, the origin of the pandemic, further supports this point. Notably, there is no high significant mutation emerged in the second phase. Together in both the phases, 26 countries have all the 9 highly significant mutations: England, USA, Wales, Australia, Scotland, Luxembourg, France, Netherlands, Iceland, Portugal, Spain, Belgium, Germany, Denmark, Greece, Sweden, China, Russia, Singapore, Taiwan, Thailand, Georgia, India, Jordan, Canada and Chile. Among the African countries, Congo carries 7 of the highly significant mutations. Most of the Asian countries possess the majority of the highly significant mutations. Though the data from the South American countries are limited, the majority of the highly significant mutations are found among all the South American countries.

**Figure 5:**
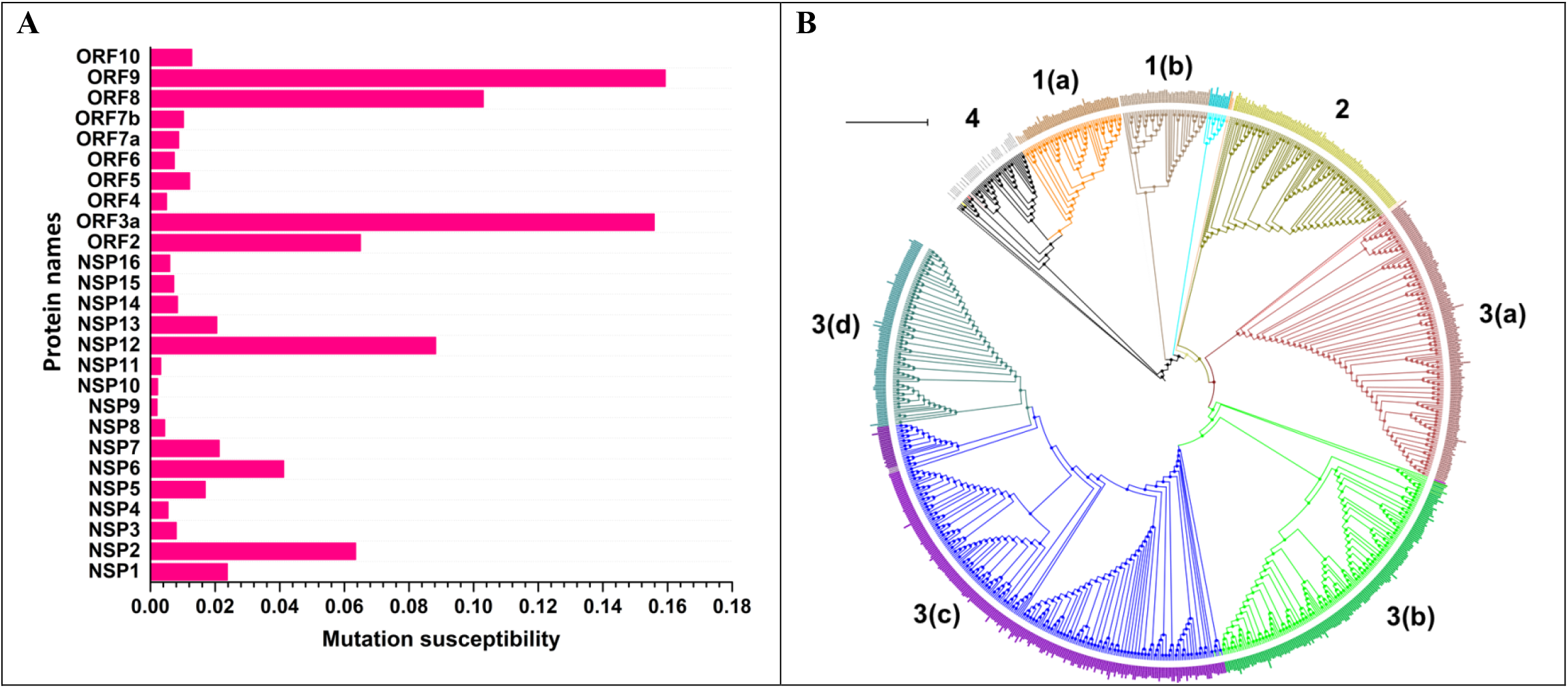
A) Bar diagram illustrating the mutation susceptibility rate (%) of 26 proteins in phase 2. Among 26 proteins, ORF9 is found to have the highest mutation susceptibility rate of ~0.16 %, followed by ORF3a (0.15585), ORF8 (0.10298), NSP12 (0.08823), ORF2 (0.06495), NSP2 (0.06345) and NSP6 (0.04118). B) Phylogram showing evolution of four different SARS-CoV-2 clades. The three major clades are indicated by different colors and are labeled as 1-3: 1(a), 1(b), 2, 3(a), 3(b), 3(c) and 3(d) correspond to G251V-ORF3a & L37F-NSP6, L37F-NSP6, L84S-ORF8, D614G-spike, P323L-NSP12, R203K-ORF9 & G204R-ORF9, D614G-spike & P323L-NSP12, D614G-spike, P323L-NSP12 & Q57H-ORF3a and D614G-spike, P323L-NSP12, Q57H-ORF3a & T85I-NSP2 mutations respectively. Note that the sub-clades are indicated by alphabets.

**Figure 6:**
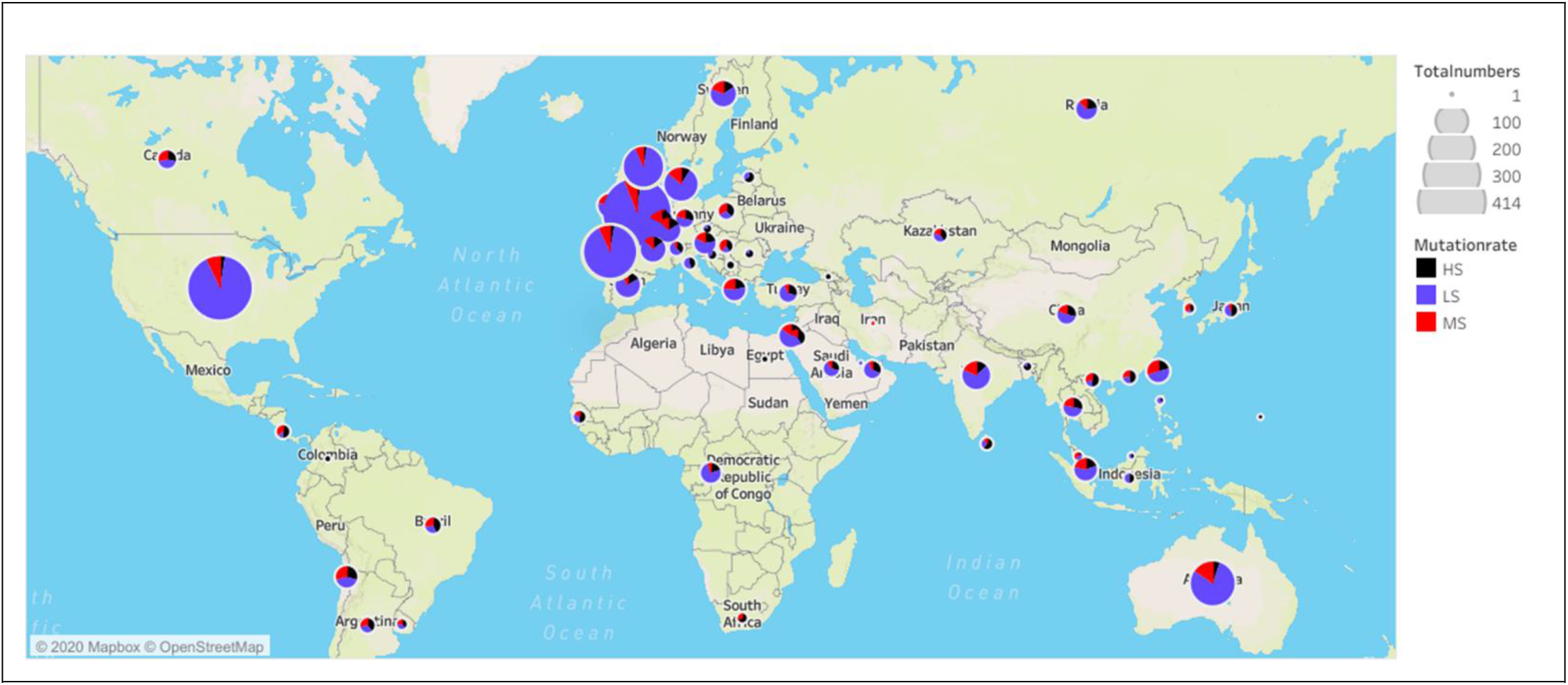
Demographic distribution of frequency of occurrence of high (black), moderate (red) and low (blue) significant mutations found in phase 2. The size of each pie chart indicates the number of sequences considered for the corresponding country. Note the presence of high significant mutation(s) in all the countries.

Further, there are 27 mutations found in phase 2 with a moderate significance (**Figure 3**), out of which, 15 are common to both the phases (**Figure 3**). The remaining 12 mutations have emerged as significant mutations in the phase 2. There are some mutations that are moderately significant in phase 1, but, absent in the phase 2 due to lack of sequence information in the latter. These mutations are found in various countries (78) and majority of the mutations (~63%) have evolved in the phase 1, indicating the positive selection pressure. In the second phase, Australia has the highest number of moderate mutations (25 out of 27) followed by USA (24) and United Kingdom (England, Wales and Scotland carry MS mutations in the range of 20 to 23) (**Figure 3**). Denmark also has 12 MS mutations. Among the Asian countries, India and Taiwan carry the highest number of moderate mutations (11). Chile has 10 of the moderate mutations. Most of the moderate mutations are absent in African countries, perhaps due to the limited availability of the data. There are 527 low significant mutations found in the second phase. Among these low significant mutations, 251 of them are common to both the phases indicating the positive selection pressure. These mutations are randomly seen across several nations. Among the 84 countries considered here in both the phases, England, USA, Wales, Australia and Scotland occupy top 5 positions with respect to the overall mutation rate.

Some notable substitutions, co-occurring mutations, deletions and insertions that occur with a low to moderate significance in phase 2 and may have a significant influence in the viral pathogenic mechanism are: co-deletion (dependent) of K141, S142 and F143, deletion of M85, co-deletion (dependent) of G82 and H83, D75E, co-occurring (dependent) S135N and Y136 deletion in NSP1, D268 deletion, G212D and co-occurring (dependent) I559V and P585S in NSP2, F10L, V198I (MS in phase 1 (1.34%), LS in phase 2 (0.7%)), P91S, T166I, H237R, T371I, S211F and G339S in NSP2 (more than 2% of among the second phase mutations), A58T,P153L and T428I in NSP3, F308Y(MS), T295I(0.3%), M33I (LS 0.19%) in NSP4, G15S and K90R (MS in phase 1 (1.14 %) and LS in phase 2 (0.1%)) in NSP5, S25L (1.5% with the highest rate in USA and Israel) and S26F (LS-0.05%) in NSP7, A97V and A449V in NSP12, A18V and P504L and Y541C co-mutation in NSP13, F233L and A320V in NSP14, V13L in ORF3a, S24L and A62L in ORF8, D936Y in spike and P13L,D103Y,S193I,S194L and S197L (each occur about 1%) in nucleoprotein. Very interestingly, a deletion in ORF8 region is found in Singapore and Taiwan. In these sequences, the ORF8 encodes only 9 amino acid peptide instead of the 121 long amino acid protein. This mutation is suggested to be associated with SARS-CoV-2 infection found in Taiwan and Middle East^66^.

Seven SARS-CoV-2 proteins (NSP2, NSP6, NSP12, spike, ORF3a, ORF8 and nucleocapsid protein) have at least one of the HS mutations and 10 proteins (NSP2, NSP3, NSP4, NSP5, NSP7, NSP13, ORF3a, membrane protein, ORF8 and nucleocapsid protein) have at least one moderate mutations. The rest of the proteins have only low significant mutations (**Figure 4**). The normalization of the mutations with respect to the protein length indicate that nucleoprotein, ORF3a and ORF8 are highly susceptible for changes and the mutation susceptibility falls in the following order in phase 2: nucleoprotein > ORF3a > ORF8 > NSP12 > spike > NSP2 > NSP6 > NSP1 > NSP7 > NSP13 > NSP5 > ORF10 > membrane protein > ORF7b > ORF7a > NSP14> NSP3 > ORF6 > NSP15 > NSP16 > NSP4 > envelope protein > NSP8 > NSP10 > NSP9 > NSP11 (**Figure 5(A)**).

The structural (**Figure 4**) and functional perspectives of the SARS-CoV-2 protein mutations reported here provide some useful information for the therapeutics. The PL-pro domain and macrodomain of NSP3 can be potential drug targets as they are comparatively resistant to mutations^67^. Occurrence of S25L and S26F in NSP7 are at the proximity of NSP8 interface, NSP8 mutations M129I (at the interface of NSP8-NSP12) and R51C/L (at the RNA interacting region) and proximity of P323L-NSP12 to the NSP8 binding site indicate a possible modulation in the viral replication mechanism. Further, majority of the amino acids in NSP6 mutates either to F or to a more hydrophobic amino acid implicating the role of this mutations in host membrane manipulations. Similarly, P134S-NSP16 found in the cavity that is in proximity to the SAM binding site may have a role in replication modulation **(Figure 4, PDB ID: 7BQ7)**. In general, except NSP12, all the replicase machinery proteins have a low mutation rate; thus, can be potential therapeutic targets. Interestingly, D614G-spike, D936Y-spike and V13L-ORF3a mutations are observed in the potential B-cell and T-cell epitope regions respectively^68^, thus, may have an influence in the host defense mechanisms. In ORF3a, many low significant mutations are observed in the ion channel (**Figure 4**). Yet another notable point is a high mutability in the SR stretch of nucleoprotein, which along with the frequent ORF8 mutations justify their role in host immune evasion^62,66^ (**Figure 4**).

Several phylogenetic analyses have been carried out to understand the evolution of SARS-COV-2^16,69–72^. A phylogram generated here using the 965 unique sequences having at least one of the significant mutations (LS, MS and HS) by considering the reference sequence (NC_045512.2) as the root indicates the evolution of 4 major clades (**Figure 4(B)**). 3 of the clades are due to the high significant mutations L37F-NSP6, L84S-ORF8 and D614G-spike, among which, the spike protein clade has the highest number of sub-clades. The fourth clade doesn’t have any high significant mutations but has moderate and low significant mutations.

Thus, the present investigation reveals the emergence of 4 major divergence from the reference sequence (Genbank ID: NC_045512.2) based on the significant mutations present in the proteome of SARS-CoV-2 for the data considered in this investigation. The comparative proteome analysis shows that out of 803 significant mutations, 531 (~66%) of them are found in the first phase and only 272 (~34%) emerged in the second phase indicating the evolution of the mutations quite early and majority of them have already been circulating around the globe. Not surprisingly, these reflect in the emergence of several clades in the phylogram. Although the variations in SARS-CoV-2 proteins reported here (based on the comparative proteome analysis) would not directly reflect the viral transmissibility, pathogenicity, virulence and immunogenicity, it clearly pinpoints that the highly significant and moderate mutations would certainly be useful in identifying an effective drug and vaccine targets that have a less heterogenicity. Indeed NSP9, NSP10, NSP11, NSP8 and envelope protein exhibit a high level of plasticity by undergoing slow mutation rate indicating them as a sensitive drug/vaccine targets. The study further would facilitate the understanding of the influence of these mutations on host viral interactions and identification of unique marker regions that could be used in the diagnostics.

## Author contribution

PPU collected and organized the whole genome sequence data of SARS-CoV-2. LPPP and CS wrote the codes to fetch the information from the data. LPPP, PPU and CS analyzed the data. LPPP, PPU, CS and TR wrote the manuscript. TR designed and supervised the entire project.

## Conflict of interest

None

## Funding

None

## Acknowledgements

The authors thank Indian Institute of Technology Hyderabad for the computational resources.

## Notes

### Competing Interest Statement

The authors have declared no competing interest.

