## Supplementary material for "Global variation in the SARS-CoV-2 proteome reveals the mutational hotspots in the drug and vaccine candidates": Methods

Kandi, Telangana-502285

† Equal contribution

†These authors contributed equally. The names are listed in the alphabetical order with respect to the last name.

**Keywords:** SARS-CoV-2 viromics, proteome analysis, mutational hotspot, mutational susceptibility, phyloproteomics, drug target, vaccine antigen, diagnostics

### Methods

SARS-CoV-2 whole genome sequences (WGS) of clinical samples collected from diverse geographical background were downloaded from the NCBI[1] (3730 sequences) and GISAID[1,2] (27659 sequences). A total of 31389 sequences were considered to analyze SARS-CoV-2 proteomic diversity across 84 countries (**Figure M1**). Subsequently, the datasets were manually curated to create a local database. The viral genome sequences with more than 15000 bases (half of the entire viral genome size) were alone considered in the creation of the database. As there is no link between the WGS deposited in the NCBI and GISAID, there is a possibility of sequence(s) duplication in the local database. The genomic sequences were then translated into individual proteins by considering the first published SARS-CoV-2 as the reference (hereafter, reference sequence) using an inhouse script. Note that if any translated region contains an undefined amino acid (due to the presence of one or more undefined nucleotides ("N") in the coding region), the region was not considered for the analysis. However, the remaining coding regions were considered for the analysis.

The country wise mutational analysis specific to each protein was analyzed through multiple sequence alignment (MSA) using CLUSTAL OMEGA [3]. For the analysis, the data was divided into two different phases. While the sequences deposited during Jan-April 17 were considered for the phase 1 analysis (10961 sequences), the sequences deposited between April 18-May17 were considered for the second phase (20428 sequences). It is worth mentioning that the data was available for 46 countries in both the phases (England, Wales, Scotland, France, Luxembourg, Netherlands, Spain, Belgium, Germany, Italy, Czech, Austria, Ireland, Denmark, Latvia, Greece, Hungary, Sweden, Switzerland, Poland, Turkey, China, Japan, Malaysia, Russia, Saudi Arabia, Singapore, South Korea, Taiwan, Hong Kong, Thailand, Vietnam, Georgia, India, Iran, Israel, USA, Canada, South Africa, Congo, Australia, Argentina, Brazil, Chile, Uruguay, Colombia). However, the data was not deposited for 22 countries in the second phase, but, available for the first phase (Iceland, Portugal, Norway, Finland, Estonia, Slovenia, Lithuania, Slovakia, Belarus, Kuwait, Nepal, Pakistan, Mexico, Ghana, Algeria, Senegal, New Zealand, Peru, Ecuador, Panama, Cambodia, Tunisia). In contrast, the data was not deposited for 16 countries in the first phase, but, available for the second (Serbia, Romania, Croatia, Philippines, Indonesia, UAE, Jordan, Qatar, Bangladesh, Brunei, Kazakhstan, Sri Lanka, Costa Rica, Guam, Gambia, Egypt).

Prior to the MSA, the 26 proteins (non-structural (NSPs 1 to 16), structural (spike, envelop, membrane and nucleocapsid) and accessory proteins (ORF3a, ORF6, ORF7a, ORF7b, oRF8 and ORF10))[4] encoded by the genomic and sub-genomic RNAs of SARS-CoV2 were isolated individually and stored country wise in the database.

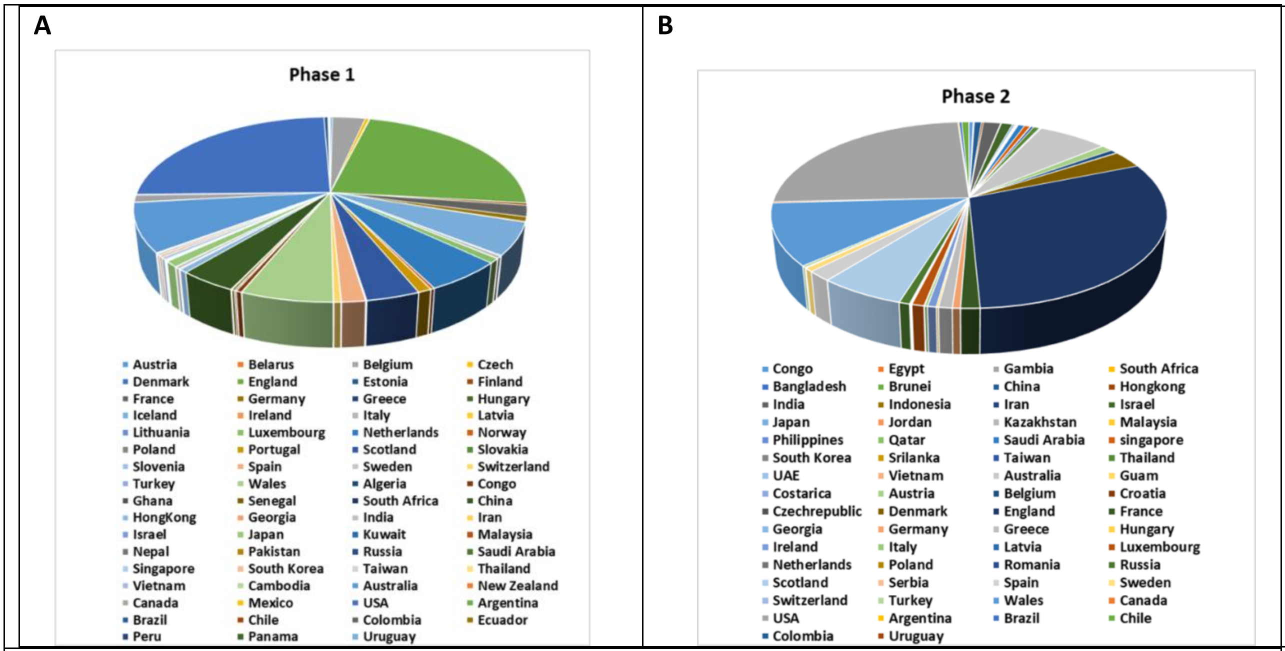


**Figure M1.** Pie chart showing the demographic statistics of SARS-CoV-2 WGS considered in the first (A) and second (B) phases.

### Calculation of the rate of mutation

The country wise MSA alignments were manually verified for each of the 26 proteins and subjected to position-wise frequency analysis of the amino acids using inhouse programs. When an amino acid at a particular position changes to a different amino acid, the number of such changes has been counted to calculate the frequency of mutation in each country. A mutation was considered as a significant (*viz.*, the mutation that does not occur by chance) only if the summation of the country wise mutation rates (**Equation 1**) of that particular mutation is at least 0.04%.

$$Rate\ of\ mutation\ (\%) = \frac{Frequency\ of\ a\ mutation\ in\ a\ country}{Total\ number\ of\ sequences} x100$$

\- Equation 1

Subsequently, the stacked bar diagrams depicting the country wise mutational rate (position wise) for each of the 26 proteins were created using Origin[5] for both the phases.

The significant mutations were further classified into three categories: highly significant (HS), moderately significant (MS) and less significant (LS). A mutation was considered as highly (dominant) or moderately significant when it occurs at the rate greater than 10% or between 1 to 10% respectively. On the other hand, a mutation was considered as less significant when it occurs at the rate between 0.04 and 1%.

Further, the mutation susceptibility rate of a protein was defined as the sum of the rate of all the mutations (**Equation 1**) that occur in that particular protein divided by its length. The overall SARS-CoV-2 mutation rate in a country was calculated by summing up the rate of mutation of all the proteins in that particular country.

The Pymol suite[6] was used to locate the mutations occurring in second phase on the 3D structures of the SAR-CoV-2 proteins wherever it is applicable.

### Construction of phyloproteomic tree

Initially, the whole genome sequences, from the second phase, that do not have undefined nucleotides ('N') were translated into 2460 single sequences with each containing 26 translated proteins (5'UTR, 3'UTR, ORF1ab stem loop 1, ORF1ab stem loop 2, ORF10 stem loop 1 and ORF10 stem loop 2 were excluded). Subsequently, 965 non-redundant protein sequences that have at least one of the 803 significant mutations (LS or MS or HS) were selected for the phylogenetic tree construction. The sequences with insignificant mutations (occurs less than 0.04%) were ignored. The selected sequences were subjected to alignment using MAFFT [7] and phyloproteomic tree was constructed using the maximum likelihood method in IQ-TREE software[8]. Itol tool[9] is used to visualize and analyze the phylogram.
